# Automatic Change Detection of Human Attractiveness: Comparing Visual and Auditory Perception

**DOI:** 10.1101/2025.10.12.681768

**Authors:** Meng Liu, Jin Gao, Werner Sommer, Weijun Li

## Abstract

Change detection of social cues across individuals plays an important role in human interaction. Here we investigated the automatic change detection of facial and vocal attractiveness in 19 female participants by recording ERPs. We adopted a ‘deviant-standard-reverse’ oddball paradigm where high- or low-attractive items were embedded as deviants in a sequence of opposite attractive standard stimuli. Both high- and low-attractive faces and voices elicited mismatch negativities (MMNs). Furthermore, low-attractive versus high-attractive items induced larger mismatch negativities in the voice condition but larger P3 amplitudes in the face condition. These data indicate that attractiveness can be automatically detected but that differences exist between facial and vocal attractiveness processing. Generally, change detection seems to work better for unattractive than attractive information possibly in line with a negativity bias.

## 1. Introduction

Detecting and coping appropriately with environmental novelty and change is highly important. In humans responding to environmental change has been investigated in oddball paradigms, where rare stimuli unpredictably occur on a background of frequent standard stimuli. These studies show that change (oddball) detection takes place not only with respect to physical stimulus features (e.g., color, line orientation, motion direction, spatial frequency, intensity or pitch) but also along more complex and socially relevant dimensions, such as posture (for a review, see Rensink, 2002). Interestingly, some studies demonstrated that unpleasant valence images can produce larger P1 amplitudes than pleasant and neutral images (for a review, see Olofsson et al., 2008). These findings are consistent with a considerable literature about a negativity bias (or negative superiority effect), suggesting that, as compared to positive stimuli, negative stimuli are processed more efficiently (for a review, see Rozin & Royzman, 2001). Physical attractiveness and unattractiveness is an important dimension of social relevance, involving aspects of facial and vocal attractiveness and plays a central role in human interaction (Hahn & Perrett, 2014; Liu et al., 2023). In the present study, we used ERP recordings to assess the automaticity of detecting facial and vocal attractiveness and whether it entails a negativity bias.

There is extensive evidence in support of the evolutionary functions of attractiveness and its role in daily life. It is generally agreed that attractiveness plays a central role in the assessment of mate value (Darwin, 1871; for a recent review, see Hahn & Perrett, 2014). The perception of attractiveness may help to select high-quality mates and transmit one’s genes to the next generations (Gallup & Frederick, 2010). However, attractiveness also plays an important role in other social contexts. Thus, attractive individuals tend to be more successful in friendship formation, job interviews, political elections and to earn more money (Wang et al., 2010; Li et al., 2024). Besides, attractive people are usually associated with positive personality traits, observed for both visual appearance and vocal properties (Rhodes, 2006; He et al., 2022; Wu et al., 2022; Niimi & Goto, 2023).

Research on attractiveness perception has focused on facial attractiveness, which can be perceived even when faces are masked or presented rapidly (Olson & Marshuetz, 2005; Shang et al., 2025) and may direct participants’ attention in the absence of consciousness (Hung et al., 2016). EEG studies show that attractiveness may affect early perceptual and cognitive processing of faces, as indicated by the N170 component, which reflects structural face representations, and by the anterior P2 component (120–220 ms), suggesting a fast attentional bias to attractive opposite-sex faces (van Hooff et al., 2011; Liu et al., 2023). Facial attractiveness also affects later cognitive processes, as reflected in the centro-parietal late positive complex (LPC), linked to motivated attention (Marzi & Viggiano, 2010; Liu et al., 2023).

Although overall attractiveness of a person substantially relates to visual cues from face and body (Yu & Shepard, 1998), it is also influenced by her or his voice (Pisanski & Feinberg, 2018). Indeed, both facial and vocal attractiveness have been linked to traits indicating sex hormone levels and health (Puts et al., 2012) and attractiveness judgments often co-vary across modalities (Hughes & Miller, 2016). Previous studies also suggested that averageness and sexual dimorphism are common factors influencing facial and vocal attractiveness judgments (Hönn & Göz, 2007). However, compared with the relatively extensive research on facial attractiveness, most studies on vocal attractiveness have been conducted in the evolutionary domain, suggesting that non-verbal voice features, such as pitch, provide ecologically relevant information about the speaker (Pisanski & Feinberg, 2018). Only a few studies have explored the neurophysiological mechanism of perceiving vocal attractiveness (Bestelmeyer et al., 2012; Hensel et al., 2015; Zhang et al., 2020; Liu et al., 2023).

For example, Zhang et al. (2020) administered an implicit tone detection task and an explicit attractiveness judgment task with short voice samples, using the ERP technique. In both tasks N1 amplitudes were larger for attractive than unattractive voices, whereas attractive voices elicited larger LPC amplitudes only in the explicit task. Hence, vocal attractiveness processing during early stages appears to be rapid and mandatory, whereas during later elaborated stages, vocal attractiveness is processed strategically and aesthetics-based, requiring attentional resources.

Of note, the research on facial and vocal attractiveness mentioned above, mainly adopted explicit tasks. Although, the perception of attractiveness should occur automatically at early stages of attention, before any significant cognitive investment is made (Zhang et al., 2018), there is little research involving pre-attentive conditions. In the ERP, fast pre-attentive change detection can be indicated by the mismatch negativity (MMN) and P3 components (Atienza et al., 2001). The auditory MMN (aMMN) with a frontocentral scalp distribution is usually observed around 100-250 ms after the onset of a deviant sound (Näätänen et al., 2007), and the visual MMN (vMMN) appears 100-300 ms after visual deviants and shows a posterior scalp distribution (Kimura et al., 2011). The parietally distributed P3 component, elicited when events require an updating of stimulus representations held in working memory, by involuntary attention capture or conscious deviance detection, usually peaks around 300 ms (Polich, 2007). The P3 may occur in implicit tasks but is usually much more pronounced in explicit tasks.

Although previous studies have found that people can automatically recognize some high-level face features (e.g., expression and gender) (Kecskés-Kovács et al., 2013; Rellecke et al., 2012), few studies have focused on the automatic processing characteristics of human attractiveness information. Using EEG, some of these studies found that face-sensitive visual areas can discriminate implicitly and rapidly between levels of facial attractiveness (Luo et al., 2019; Rellecke et al., 2011) and that attractive faces can be processed automatically by both males and females (Zhang et al., 2018). However, other research indicates that facial attractiveness processing is not mandatory since attractiveness-related modulations of brain responses were only a trend during a gender decision task (Schacht et al., 2008). Hence, there may be early automatic processes of attractiveness detection, and later controlled processes. It is even less clear whether vocal attractiveness information can be perceived automatically and how this perception compares to facial attractiveness.

A further question of interest is the contrast between high and low attractiveness and how it depends on the facial versus vocal domain. Interestingly, some studies demonstrated that change detection of emotional valence is more sensitive for negative than positive valence (Todd et al., 2023; Kreegipuu et al., 2013). These findings are consistent with a considerable literature about a negativity bias (or negative superiority effect), suggesting that, as compared to positive stimuli, negative stimuli are processed more efficiently (for a review, see Rozin & Royzman, 2001). This might hold also for unattractiveness as compared to attractiveness.

The present study adopted a deviant-standard reverse oddball paradigm (Conde et al., 2015) in which probability and attractiveness were manipulated in order to investigate the correlates of implicit facial and vocal attractiveness processing. Our prominent aim was to investigate whether change detection during implicit facial and vocal attractiveness perception is modulated by high versus low attractiveness categories. Specifically, given that automatic change detection processes are facilitated by the emotional quality of faces (Susac et al., 2003) or vocalizations (Schirmer & Kotz, 2006), we hypothesized that also attractiveness differences would induce change detection processes as indexed by the MMN and P3 components. If change detection is facilitated by unattractive faces or voices (i.e., unattractive stimuli associated with avoidance), we expected increased MMN and P3 amplitudes for unattractive deviants relative to attractive standard stimuli. Also, if change detection is facilitated by attractive cues (i.e., attractive stimuli associated with pleasure and approach), one should expected increased MMN and P3 amplitudes for attractive deviants compared with unattractive standard stimuli. In line with the negativity bias, we expect the effect for unattractive deviants to be more pronounced than for attractive deviants.

## 2. Methods

### 2.1 Participants

Twenty-one female undergraduate students participated in the experiment for reimbursement. All of them reported normal or corrected-to-normal vision, no history of hearing or neurological impairments, and to be right-handed. This study was approved by the Ethics Committee of Liaoning Normal University. Each participant had given informed consent. Owing to excessive EEG artifacts, data from two participants were excluded; the final sample included 19 female participants, aged 18 to 24 years (*M* = 21.62 ± 1.3 years). In order to reduce complexity we tested only female participants with stimuli from male actors.

### 2.2 Materials

The initial pool of facial stimuli consisted of pictures of 30 attractive and 30 unattractive male faces that had been used in a previous study (Yang et al., 2015). All faces showed neutral expressions, were taken from a frontal view and with eye gaze directed at the observer. To exclude non-facial information, external features (i.e., hair, ears) were removed and images were cropped to an approximate size of 260×300 pixels. Forty undergraduate students who did not participate in the ERP experiment validated the 60 faces on a 7-point rating scale (ranging from 1 = very unattractive to 7 = very attractive). The five most attractive (*M* = 5.28) and five most unattractive (*M* = 2.01) faces were selected as stimuli for the ERP experiment. Besides, each of these ten faces was also set with a red dot on the nose as target stimuli for the experimental task.

The initial pool of male vocal stimuli consisted of the 40 attractive and 40 unattractive vowel syllables from a previous study (Zhang et al., 2020). These syllables, recorded with Adobe Audition software and digitized at 16-bit/44.1kHz, were provided by professional male speakers in an emotionally neutral prosody. Praat software (http://www.fon.hum.uva.nl/praat/) was used to equalize the duration (400 ms) and normalize the intensity (70 dB) of the syllables. Thirty further undergraduate students rated the vocal materials on a 7-point scale (ranging from 1 = very unattractive to 7 = very attractive). From the 40 rated syllables the five most attractive (*M* = 5.09) and five most unattractive (*M* = 2.74) ones were selected as stimuli for the ERP experiment. Furthermore, each of the ten syllables was also mixed with white noise, serving as target stimuli.

### 2.3 Procedure

Participants were comfortably seated in an electrically shielded room at a distance of 90 cm from a computer screen. The deviant-standard reverse oddball paradigm was conducted in separate face and voice conditions (see Figure 1), with the order of conditions counterbalanced across participants. Visual stimuli were presented at the center of the screen, and auditory stimuli were presented via headphone at a comfortable listening level. In both conditions, simultaneous appearance of a fixation cross and warning tone (500 ms) at the beginning of each block was followed by a sequence of face or voice stimuli. The duration of each face/voice stimulus was 400 ms, and the inter-trial-intervals (ITI) varied between 1.3 and 1.5 s (see Figure 1). Participants were instructed to complete an attractiveness-unrelated task by pressing the space key as quickly as possible whenever they saw a face with red nose (targets) in the face condition or heard a voice mixed with white noise (targets) in the voice condition. Attractive and unattractive stimuli were presented as both standards and deviants to eliminate the differences in the physical properties between faces/voices. Overall, the ERP experiment was composed of four conditions, (1) attractive face standards and unattractive face deviants, (2) unattractive face standards and attractive face deviants, (3) attractive voice standards and unattractive voice deviants, and (4) unattractive voice standards and attractive voice deviants. Each of these four conditions contained 600 trials in total, including 450 standards (75%), 90 deviants (15%) and 60 targets (10%). The 600 trials of each condition were separated into three blocks of 200 trials each, yielding a total of 12 blocks for the whole experiment.

**Figure 1.**
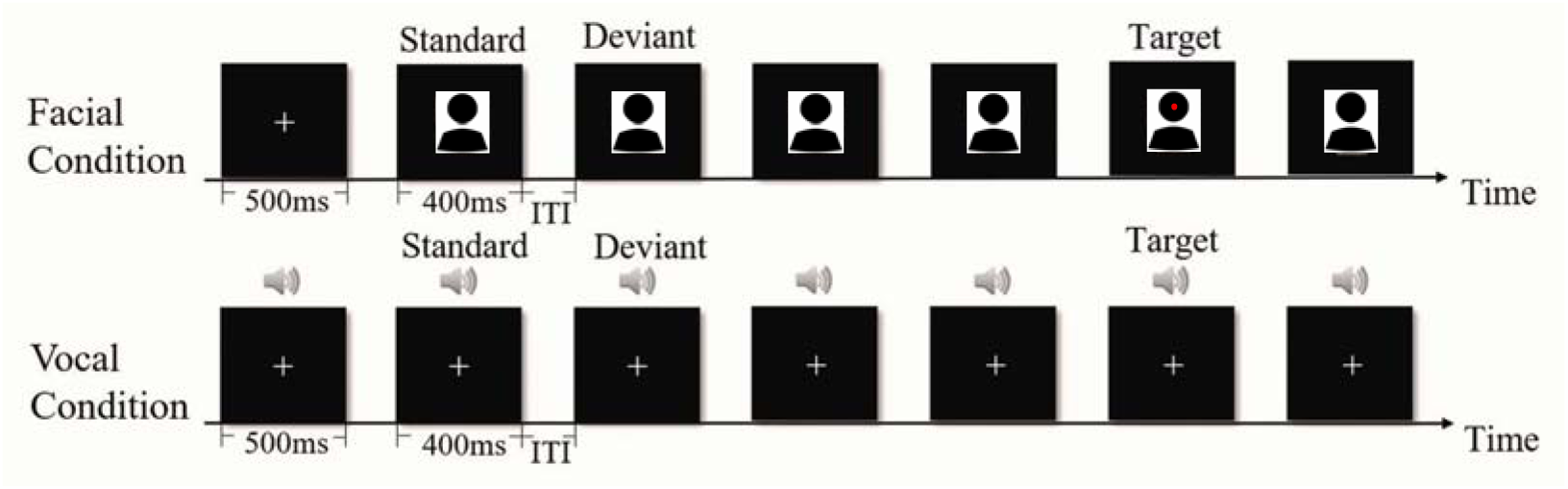
Schematic illustration of the unattractive in attractive Oddball condition for face condition and voice condition. Stimulus duration was 400 ms for both faces and voices; the inter-trial-intervals (ITI) varied between 1300-1500 ms. Note: To avoid the inclusion of photographs and any other identifying information of people, human icon was substituted to illustrate the relevant concept.

All trials within each of the four conditions were pseudorandomized and there were at least three standards in-between two deviants. Targets (faces with red noses or voices with white noise) were always derived from standards and randomly mixed into all trials to ensure that participants attended to the stimuli. In addition, the 12 blocks were arranged in four sequences to counterbalance experimental conditions and order of presentation. Each participant completed one of the four block sequences and each condition began with 10 practice trials. The entire duration of the experiment was about 2.5 h, including EEG preparation, practice and experiment proper; participants rested for 1-3 min after each block.

### 2.4 EEG recording and pre-processing

The EEG was recorded from 64 cap-mounted Ag/AgCl electrodes (ANT Neuro EEGO Inc., Germany) according to the modified 10-20 system, with CPz as on-line reference, at a sampling rate of 500 Hz. Electrode impedance was kept below 5 kΩ. During recordings, a 100 Hz low-pass filter was applied. Offline, EEG signals were re-referenced to averaged mastoids, and submitted to a 0.01-30 Hz band-pass filter. Ocular artifact correction was performed through independent component analysis in EEGLAB (Bayazıt et al., 2009). Continuous data was segmented into stimulus-locked −200 to 800 ms epochs. Signals exceeding ±80 μV in any given epoch were automatically excluded. Other artifacts were removed according to visual inspection. All trials that immediately followed a deviant stimulus were discarded from analysis.

### 2.5 Statistical analyses

We analyzed the vMMN for the face conditions, the aMMN for the voice conditions and the P3 for all conditions. The vMMN was maximal over the parieto-occipital scalp in the 200-300 ms range and was measured at left and right parieto-occipital regions of interest (ROIs; electrodes P5, P7, PO5, PO7 and P6, P8, PO6, PO8). The aMMN was maximal over the frontal-central scalp in the 200-300 ms range and measured at left and right fronto-central ROIs (F1, F3, C1, C3 and F2, F4, C2, C4). The P3 was maximal over the parietal scalp in two consecutive 150 ms time windows (350 to 650 ms) and measured at left and right parietal ROIs (CP1, CP3, P1, P3 and CP2, CP4, P2, P4). For purposes of analysis, signals for the electrodes of a given ROI were averaged. For each stimulus modality three-way repeated measures ANOVAs were applied to the mean amplitudes with Attractiveness (attractive, unattractive), Deviance (standard, deviant), and Hemisphere (left, right) as factors. Only adjusted *p*-values will be reported.

## 3. Results

### 3.1 Behavioral results

Participants performed with high accuracy in detecting the target stimuli (*M* = 94%, *SD* = 2%). Mean reaction time was 526 ms (*SD* = 75 ms). There was no difference in performance across conditions.

### 3.2 ERP results

#### 3.2.1 MMN effect for face and voice conditions

Figures 2 and 3 illustrate the MMN effect elicited by attractive and unattractive stimuli for the face and voice conditions. In the face condition, ANOVA on mean amplitudes (200-300 ms) showed that deviants elicited larger parieto-occipital negativities than standards, *F* (1, 18) = 8.25, *p* < 0.01, *η^2^* = 0.31. Besides, the interaction between Deviance and Hemisphere was significant, *F* (1, 18) = 10.17, *p* < 0.01, *η^2^* = 0.36. Simple effect analysis indicated that deviants elicited larger negativities than standards in the right hemisphere, *F* (1, 18) = 10.48, *p* < 0.01, *η^2^* = 0.37, but not in the left hemisphere, *p =* 0.09 (see Figure 3a).

**Figure 2.**
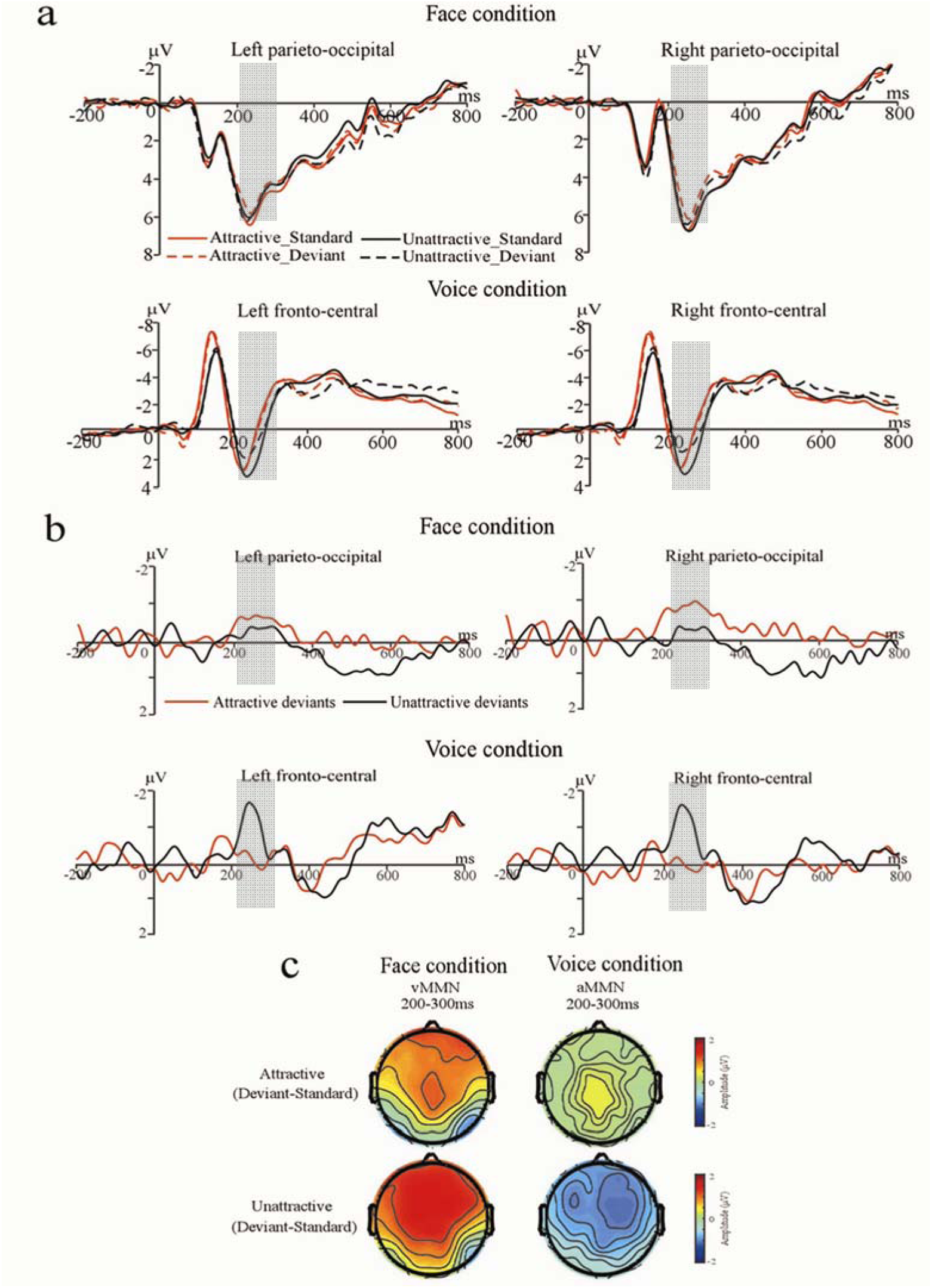
ERP results for mismatch negativities (MMN), marked by shadings. (a) ERPs elicited by standards and deviants in the parieto-occipital region (P5, P6, P7, P8, PO5, PO6, PO7 and PO8) in the face condition, and in the fronto-central region (F1, F2, F3, F4, C1, C2, C3 and C4) in the voice condition. (b) Difference waveforms (MMN) obtained by subtracting standards from deviants in each condition. (c) Topographies of ERP differences (MMN) for attractive and unattractive oddball effects in each condition.

**Figure 3.**
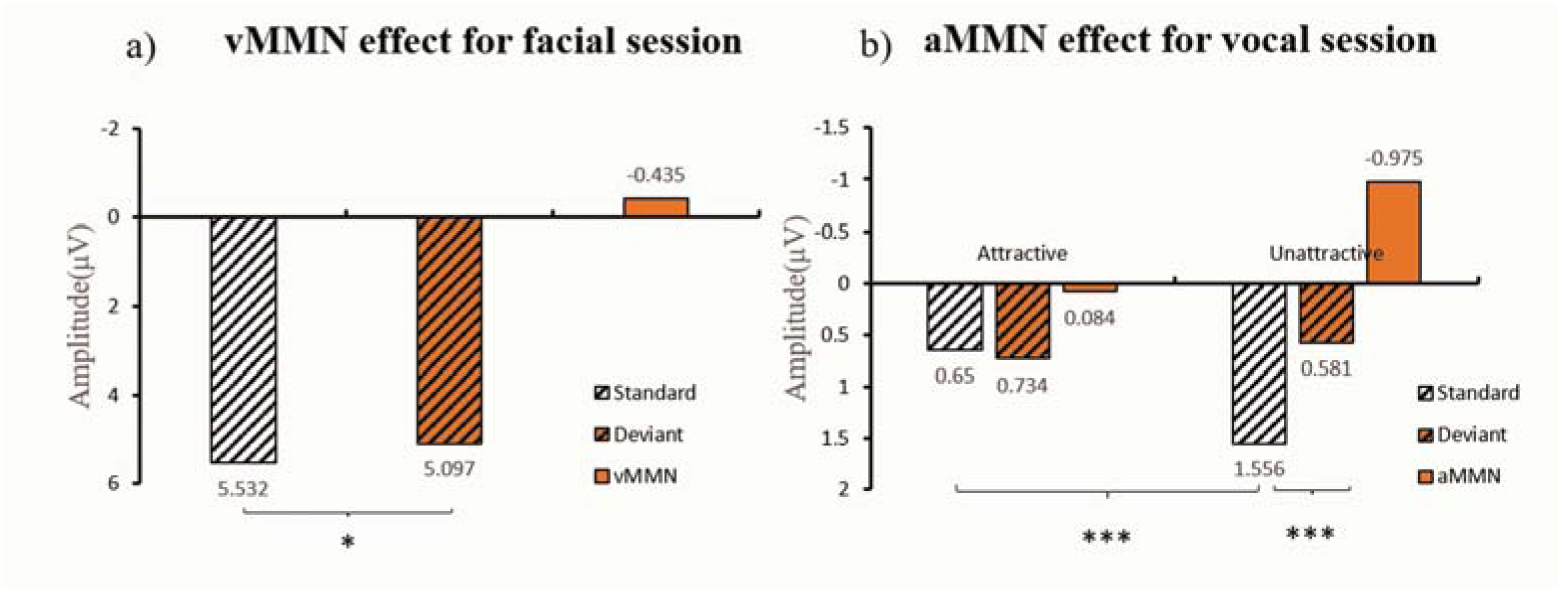
Mean amplitudes during the MMN intervals (200-300 ms). (a) Face condition: vMMN = visual mismatch negativity. (b) Voice condition: aMMN = auditory mismatch negativity. Note: \**p* < 0.05, \*\*\**p* < 0.001.

In the voice condition, ANOVA showed a main effect of Attractiveness, *F* (1, 18) = 4.05, *p* < 0.05, *η^2^* = 0.21; attractive voices elicited larger frontal-central negativities than unattractive voices. Besides, the interaction between Deviance and Attractiveness was significant, *F* (1, 18) = 10.04, *p* < 0.01, *η^2^* = 0.36. Simple effect analysis revealed that unattractive deviants elicited larger negativities than standards, *F* (1, 18) = 15.00, *p* < 0.001, *η^2^* = 0.46, but not attractive deviants, *p =* 0.81 (see Figure 3b). This interaction also revealed more negative amplitudes for attractive than unattractive standards, *F* (1, 18) = 25.53, *p* < 0.001, *η^2^* = 0.59.

#### 3.2.2 P3 effect for face and voice conditions

Figures 4 and 5 show the ERP waveforms for the P3 ROIs elicited by attractive and unattractive stimuli for the face and voice conditions. In both conditions, the P3 was analyzed in two consecutive 150 ms time windows (350 to 500 and 500 to 650 ms). In the face condition, deviants elicited larger amplitudes than standards in both the early P3 segment (350-500 ms)*, F* (1, 18) = 30.12, *p* < 0.001, *η^2^* = 0.63, and the late P3 segment (500-650 ms), *F* (1, 18) = 26.22, *p* < 0.001, *η^2^* = 0.59. Besides, for the late P3 segment the interaction between Deviance and Attractiveness was a trend, *F* (1, 18) = 4.05, *p* = 0.05, *η^2^* = 0.18. Simple effect analysis showed both, more positive amplitudes for attractive deviants relative to unattractive standards, *F* (1, 18) = 4.61, *p* < 0.05, *η^2^* = 0.21, as well as more positive amplitudes for unattractive deviants relative to attractive standards, *F* (1, 18) = 24.84, *p* < 0.001, *η^2^* = 0.59 (see Figure 5b). This interaction also revealed larger late positivities to attractive relative to unattractive standards, *F* (1, 18) = 6.82, *p* < 0.05, *η^2^* = 0.28, but no such difference for the deviants, *p =* 0.30.

**Figure 4.**
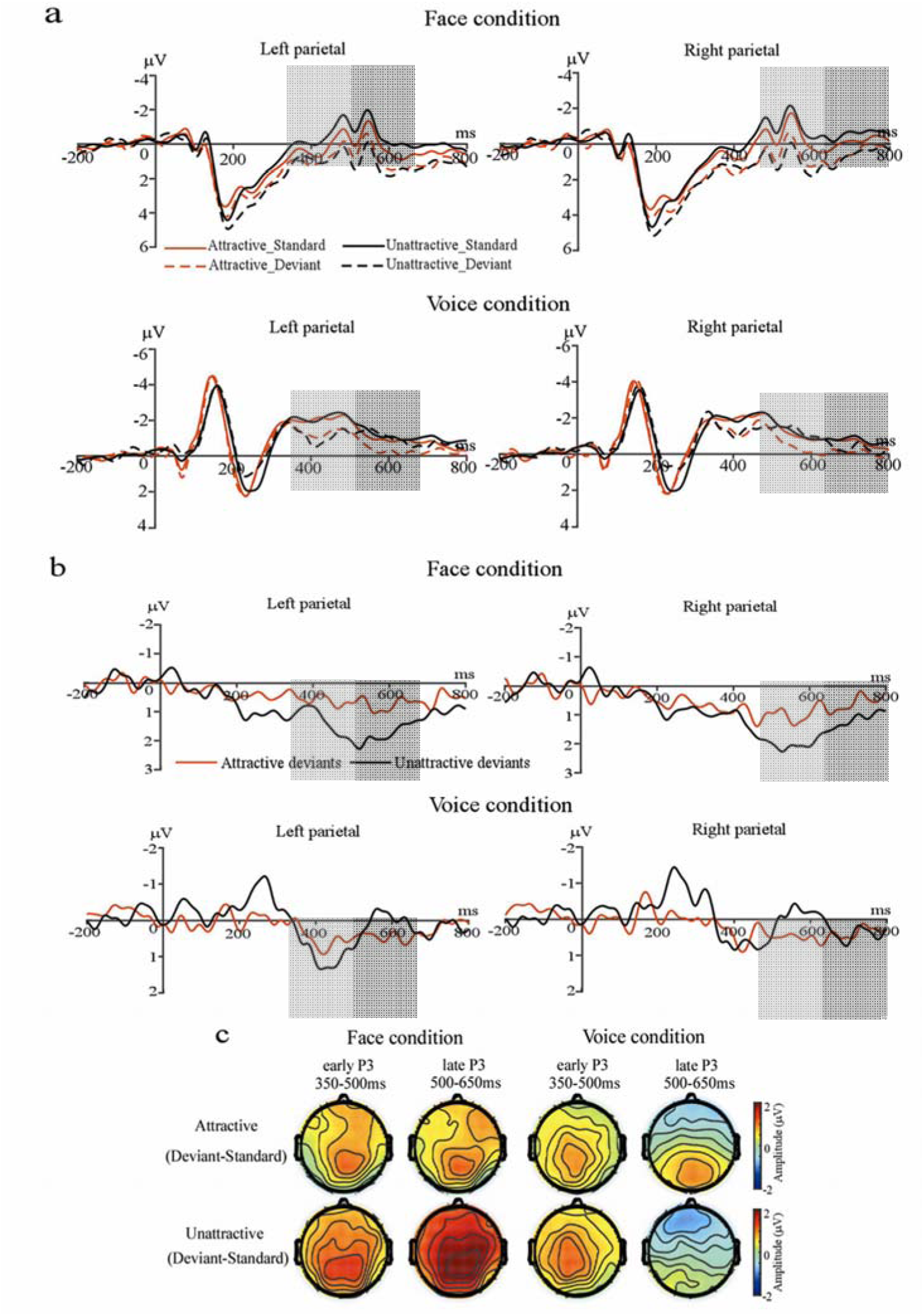
ERP results for two P3 intervals (marked by shadows). (a) ERPs elicited by standards and deviants over the parietal region (CP1, CP2, CP3, CP4, P1, P2, P3, and P4) in both face and voice conditions where the P3 was maximal. (b) Difference waveforms (P3) obtained by subtracting standards from deviants in each condition. (c) Topographies of ERP differences (P3) for attractive and unattractive oddball effects in each condition.

**Figure 5.**
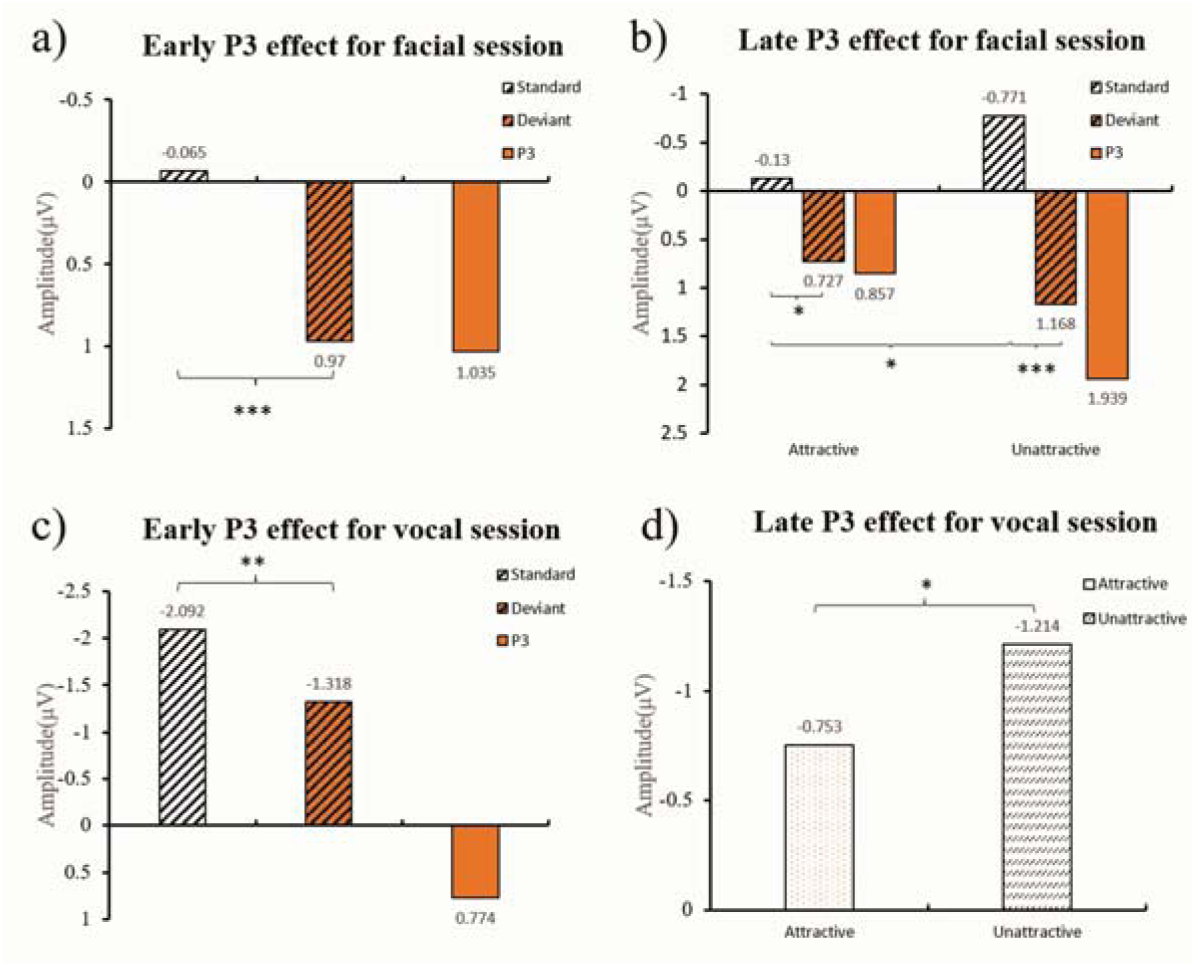
Mean P3 amplitudes. (a and b): Face conditions, early (350-500 ms) and late (500-650 ms) segment, respectively. (c and d): Voice conditions, early (350-500ms) and late (500-650 ms) segment, respectively. Note: \**p* < 0.05, \*\**p* < 0.01, \*\*\**p* < 0.001.

In the voice condition, statistical analysis of the early P3 segment (350-500 ms) revealed larger positivities to deviants than to standards*, F* (1, 18) = 9.38, *p* < 0.01, *η^2^* = 0.34. Besides, the interaction between Deviance and Hemisphere was significant, *F* (1, 18) = 4.65, *p* < 0.05, *η^2^* = 0.21. Simple effect analysis indicated that deviants elicited larger early positivities than standards in both the left hemisphere, *F* (1, 18) = 12.65, *p* < 0.01, *η^2^* = 0.41, and the right hemisphere, *F* (1, 18) = 6.07, *p* < 0.05, *η^2^* = 0.25. For the late P3 segment (500-650 ms), the main effect of Attractiveness was significant, *F* (1, 18) = 5.47, *p* < 0.05, *η^2^* = 0.24, with attractive voices eliciting larger P3 amplitudes than unattractive voices (see Figure 5d).

## 4. Discussion

The present study investigated the automatic detection of facial and vocal attractiveness by assessing passive change detection for faces and voices in ‘deviant-standard-reverse’ oddball paradigms. We present ERP evidence that attractiveness information in general, but especially unattractiveness, is detected automatically. We also demonstrate that automatic processing of facial and vocal attractiveness shows both similarities and discrepancies. Specifically, compared to high-attractive information, low-attractive deviance information induced a larger MMN effect for voices but a larger P3 effect in faces.

### 4.1 The change detection of facial attractiveness

The MMN is thought to reflect the automatic detection of deviance between sensory input and predictive memory representations generated by repeated (standard) stimuli (Czigler, 2007). In the present study, both attractive and unattractive faces elicited ERP components distributed over parieto-occipital regions, consistent with classic vMMN findings (Kimura et al., 2011). This observation indicates that sequentially presented attractiveness information (standard stimuli) is stored in a memory representation that facilitates predictions about what will happen next (Pinheiro et al., 2017). When the visual input (deviant stimuli) does not match the top-down prediction, a prediction error to the unexpected change of attractiveness reflected in the vMMN component was elicited. Since both attractive and unattractive faces elicited vMMN effects, the present experiment indicates that participants are highly sensitive to facial attractiveness changes in both directions.

Although attractive and unattractive deviant faces enhanced the P3 component, the effect was larger for unattractive deviants. The observation of a bias towards unattractiveness seems consistent with the large body of evidence about a negativity bias in many domains such as emotional experience, impression formation and intimacy (for a review, see Rozin & Royzman, 2001). Unattractive faces, which are considered to signal bad health status and/or low immunocompetence, may cause avoidance motivation. In contrast, attractive, presumably healthier faces may induce approach motivation (Grammer et al., 2001). From an evolutionary perspective, when it comes to mate choice, the key to maximize the odds for survival and reproduction may not simply be to associate with fit individuals but also to avoid unhealthy individuals (Baumeister et al., 2001). In the present study, when participants were dealing with changes of attractiveness, their motives to avoid unattractive faces may have taken precedence over their motives to approach attractive faces. According to Elliot (2013), avoidant motives can focus attention on negative rather than positive situations and options, and may take precedence over other goals and considerations. Meanwhile, the P3 component is concomitant with a change or updating of the stimulus representations governed by attentional processes (Polich, 2007). Thus, the motives to avoid unattractive faces may have directed attentional processes more powerfully and led to stronger P3 effects in the present study than the motives to approach attractive faces.

### 4.2 The change detection of vocal attractiveness

In contrast to the symmetric pattern of vMMN elicited by deviant faces, a bias for unattractive deviants was observed in the aMMN for voices. This observation may relate to differences between perceptual representations in the auditory and visual systems. Humans characterize concrete objects (e.g., location and shape) by relying on visual perception, but “abstract objects” (e.g., melodies and sound patterns) may be characterized better in the auditory domain (Winkler et al., 2009; Winkler & Czigler, 2012). Besides, according to the model of Auditory Scene Analysis (ASA) (Smoliar & Bregman, 1991), aMMN elicitation can correspond to individuals’ active exploration of alternative interpretations of the input (conveyed by top-down biasing) (Winkler et al., 2009). Consequently, in the present study, the superiority of processing ‘abstract patterns (in a ‘standard-deviant’ way) of attractiveness’ in the auditory system may cause an early-stage negativity bias, indexed by the aMMN. Alternatively, unattractive voices (e.g., a rough timbre), which are characterized by relatively simple acoustic properties as compared to attractive voices (Omori et al., 1997), may be available rather early during auditory processing and elicit an aMMN effect. in contrast, attractive voices (e.g., smooth voice) may not stand out as saliently from an unattractive background. In contrast, when it comes to faces, attractive and unattractive facial properties may benefit from their relatively affluent information (Rezlescu et al., 2015) and may be similarly distinguishable on a standard background of opposite attractiveness.

Interestingly, unlike the bias for unattractive deviant faces, the bias for unattractive deviant voices was not reflected in the P3 component. Different degrees of familiarity for facial and vocal attractiveness may help to explain this observation. According to Rezlescu and colleagues (2015), human attractiveness impressions usually rely more on faces than voices. In other words, people may be more familiar with facial attractiveness. Thus, they may more easily assign intrinsic significance to facial attractiveness than to vocal attractiveness. Analogously, in the present study, the better familiarity with facial than vocal attractiveness of syllables may have elicited a larger P3 effect for the former during cognitive stimulus evaluation.

### 4.3 Limitations and future directions

As a result of our investigations, suggestions may be made for future research. Firstly, researchers could utilize other approaches (i.e., the equiprobable paradigm) to control refractory effects and generalize the present findings. Secondly, distinctiveness may modulate the negativity bias observed in the vocal task. According to (Zäske et al., 2018), ‘voice ratings based on vowels exhibited a moderate but significant negative correlation between attractiveness and distinctiveness,’ the bias to unattractive voices may not only be induced by its lower attractiveness but also by its higher distinctiveness in the present study. Although we successfully manipulated different levels of vocal attractiveness (evidence: the main effect of attractiveness showing that attractive voices induced larger negativities than unattractive ones was observed), the potential role of distinctiveness is worthwhile for further exploration. Thirdly, change detection of attractiveness can extend to test male participants and focus on potential gender differences.

## 5. Conclusion

Voices or faces, deviating in attractiveness were robustly detected automatically but differing in time course, depending on valence and modality. Specifically, the detecting attractive items was weaker than detecting unattractive ones, demonstrating a negativity bias; this bias was present in the early-stage processing voices and in late stages of processing faces. We speculate that – at least in females, as tested here – this negativity bias in (un)attractiveness detection may serve the greater importance to avoid social partners with unfavorable health or immune status than to approach those with apparently good status.

## Author contributions

**Meng Liu:** Conceptualization, Data curation, Formal analysis, Software, Writing - original draft, Writing - review & editing. **Jin Gao:** Writing - original draft, Writing - review & editing. **Werner Sommer:** Conceptualization, Supervision, Validation, Writing - review & editing. **Weijun Li:** Funding acquisition, Supervision, Writing - original draft, Writing - review & editing, Project Administration.

## Data availability statement

The data presented in this study are available on request from the corresponding author.

